# Can we predict which species win when new habitat becomes available?

**DOI:** 10.1101/562959

**Authors:** Miki Nomura, Ralf Ohlemüller, William G. Lee, Kelvin M. Lloyd, Barbara J. Anderson

**Affiliations:** Department of Geography, University of Otago, PO Box 56, Dunedin, New Zealand; Manaaki Whenua Landcare Research, Private bag 1930, Dunedin, New Zealand

## Abstract

Land cover change is a key component of anthropogenic global environmental change, contributing to changes in environmental conditions of habitats. These changes can lead to the redistribution of species and shifts in the functional composition and properties of ecosystems. Deforestation is globally the most widespread anthropogenically driven land cover change leading to conversion from closed forest to open non-forest habitat. The consequences of these functional habitat changes on species distributions are only poorly understood. This study investigates the relative roles of geographic features, species climatic niche characteristics and species traits in determining the ability of open-habitat plant species to take advantage of recently opened habitats. We use current occurrence records of 18 herbaceous, predominantly open-habitat species of the genus *Acaena* (*Rosaceae*) to determine their prevalence in recently opened habitat. Geographic features of the spatial distribution of open habitat, species’ climatic niche characteristics, and species traits related to dispersal were tested their correlation with species’ prevalence in anthropogenically opened habitat. While primary open habitat (naturally open) was characterised by cold climates, secondary open habitat (naturally closed but anthropogenically opened) is characterised by warmer and wetter conditions. We found high levels of variation in the prevalence of secondary open habitat among the investigated species indicating differences between species in their ability to colonise newly opened habitat. For the species investigated, geographical and climatic niche factors showed generally stronger relationships with species’ prevalence in secondary open habitat than functional traits did. For small herbaceous species, geographical and environmental factors appear to be more important than species functional traits for facilitating expansion into secondary open habitats. Our results suggested that the land cover change might have triggered the shifts of factors controlling open-habitat plant distributions from the competition with forest trees to current environmental constraints.

## Introduction

Over three quarters of the global land surface have been modified by human activity [1]. In the last two decades alone, c. one-tenth (3.3 million square km) of global wilderness areas was lost [2]. Such anthropogenic land cover change affects biodiversity loss from habitat declines, and therefore can lead to the functions and distributions of species and ecosystems [3–5]. As the original (or natural) vegetation and physical properties of an area are modified, the available habitat to species and the environmental conditions will change and affect which species and ecosystems are found in that area [6].

Deforestation is a typical example of anthropogenic land cover change and, at its most basic level, results in a change from forest habitat to more open, non-forest habitat, usually scrubland or grassland. Deforestation occurred in many parts of the world following human settlement (e.g. North America [7], Europe [8] and New Zealand [9]) and is ongoing; 2.3 million square kilometres forest was lost globally between 2000 and 2012 [10]. Species distributions are strongly dependent on the environmental conditions that define habitat, and therefore, species are susceptible to land cover change [11–13]. Understanding how species respond to habitat change is important for predicting how ongoing anthropogenic land cover change may influence future species assemblages. Here, we investigate the relative contribution of landscape structure, species climatic niches and species functional traits to species’ expansion into recently opened habitats.

The effects of land cover change history on plant distributions have been reported widely in the world [14–16]. Although the history since pre-human times is not generally available, New Zealand offers good records of the land use change history since the first human settlement, because a human being settled in the land much later (c. 800 years ago) than other regions in the world. Habitats which have been available for organisms before and after anthropogenic activities, primary habitats, and those which became available after anthropogenic activities, secondary habitats, have different ecosystems. For example, a primary forest in tropical zones showed marked differences in community structure and composition from secondary and plantation forests [3]. Therefore, the expansion of secondary open-habitat following human arrival provides a new ecological opportunity for open habitat species to expand their range across these recently deforested areas.

In this study, we investigate the geographical distribution and climatic niches of 18 herbaceous species in relation to both primary and secondary open habitat in New Zealand. We assess the relative prevalence of the species in these habitats and determine the importance of three sets of factors – the geographic features, the species’ climatic niches and the species’ dispersal traits for expansion into the secondary habitats. Specifically, we address three questions;

1. What are the climatic characteristics of primary and secondary open habitats occupied by the species?
2. What are the current spatial distributions of the species in primary vs. secondary open habitat?
3. What is the relative importance of geographic features of habitat, the species’ climatic niches and species dispersal traits for expansion into secondary open habitat?

## Material and Methods

### Study Species

Occurrence records - We used occurrence records and trait data for 18 of 21 species of the genus *Acaena* occurring in New Zealand (Table S1). Three species were not used in this study because of the small number of occurrence records (< 5). The genus *Acaena* is a characteristic herbaceous element of open habitats in New Zealand, with a wide geographical and environmental range [17]. The genus is confined mostly to the southern hemisphere and comprises approximately 50 species [18, 19]. Indigenous New Zealand species of *Acaena* are prostrate, long-lived perennials, representing two main divisions based on contrasting dispersal features; the presence/absence of barbed spines on their fruits [17]. Of the 18 species selected, 17 species are native to New Zealand and one species (*A. agnipila*) is introduced from Australia and naturalised [20]. Occurrence records of these species were compiled from personal observation, surveys and reports (See a reference list in Appendix for detailed source information) and location information from online databases; New Zealand Virtual Herbarium (http://www.virtualherbarium.org.nz) and New Zealand National Vegetation Survey (https://nvs.landcareresearch.co.nz).

### Pre-human and current land cover data

New Zealand’s pre-human land cover was derived from modelled spatial data of potential suitability of New Zealand’s key forest tree species at 100 m grid resolution [21]. Current land cover was derived from the latest version of the New Zealand land cover polygon data, ‘LCDB4’ [22]. We converted pre-human and current land cover and a digital elevation model for the area [23] to rasters on 1km grid resolution using the majority rule in ArcGIS 10.2 [24].

In both land cover datasets, land cover classes were amalgamated so that each 1 km grid cell was assigned to one of three land cover types:

1. Native forest: Grid cells with any type of indigenous forest.
2. Non-forest: Grid cells with non-forest, open land cover classes, which are potentially suitable for *Acaena* species, e.g. grasslands, shrublands and gravel areas. These non-forest grid cells are here referred to as ‘open’ habitat.
3. Others: Grid cells with land cover classes that are typically not potential habitats for *Acaena* species e.g. urban area and waterbodies.

For a full list of class conversions from LCDB land cover classes into the three land cover types used in this study, see Table S2. In addition, the grid cells of current non-forest were assigned levels of openness, “high” or “low” (Table S2b).

In order to quantify the change from forest to open habitat, each 1 km grid cell was assigned one of the following three categories:

I. <u>Primary open habitat:</u> Grid cells that continuously had open habitat, i.e. are non-forest land cover in pre-human and current times.
II. <u>Secondary open habitat:</u> Grid cells that only had open habitat since human arrival, i.e. had forest land cover in pre-human times and non-forest land cover currently.
III. <u>Others:</u> Grid cells that are neither primary nor secondary open habitat. Hereafter, we refer to species occurrence records in primary/secondary open area as “primary/ secondary open occurrence records”.

Our principle metric is “species prevalence in secondary open habitat”, which quantifies effects of anthropogenic land cover change on open habitat species. We calculated the species prevalence in secondary open habitat as:

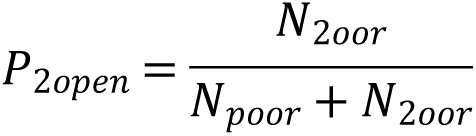

where *P*_*2open*_ is the proportion of secondary open habitat, *N*_*2oor*_ is the number of secondary open occurrence records and *N*_*poor*_ is the number of primary open occurrence records.

High values reflect that the species has a proportionally high prevalence in secondary open habitat, which we interpret as high ability to utilise newly opened habitat.

### Current climatic conditions and *Acaena* species climatic niches

Gridded average climate data (1960 - 1990) was retrieved from http://www.worldclim.org/current for four climate variables to quantify climatic conditions available in New Zealand and species climatic niches: annual mean temperature, minimum temperature of coldest month, annual precipitation and precipitation seasonality [25].

Environmental analyses were limited to climatic factors, as temperature and precipitation are likely to be primary driving factors of *Acaena* species distributions at this national spatial scale [26]. To capture the multi-dimensional climate space, an ordination, Principal Component Analysis (PCA) [27], was performed on the four climate variables using the package “stats” in R [28]. The first two ordination axes explained 61.6% and 24.0% of the variation in the climate data respectively and were here used to delineate New Zealand climate space and the *Acaena* species’ climatic niches. Hereafter, the first ordination axis is referred to as the “temperature axis” because it is strongly correlated with temperature variables and the second axis is referred to as “precipitation axis”. High values on the temperature axis indicate a cold environment, while high values on the precipitation axis indicate a dry environment.

### Correlates of prevalence in secondary open habitat

We investigated the relative importance of species geographical, environmental and functional trait features for facilitating species to move into new open habitat as it became available following human settlement. The relationship between “species prevalence in secondary open habitat” (response variable) defined above and the following indices (predictor variables) from the three main groups (Table 1) was tested with a generalized linear model using the R package “stats” with a normal error function and an identity link.

**Table 1:** Correlates of prevalence for secondary open habitat in the investigated *Acaena species*.

| Groups of factors | Factors tested | Coefficients | SE | p-values |
| --- | --- | --- | --- | --- |
|  | Intercept | 0.85 | 0.39 | 0.06 |
| Geography | Species' current range size | -0.48 | 0.41 | 0.28 |
|  | Preference for open habitat | 0.83 | 0.90 | 0.39 |
|  | Availability of secondary open habitat | 1.84 | 2.20 | 0.43 |
| | Mean elevation of current range | $-4.2 \times 10^{-4}$ | <0.01 | 0.43 |
| Climatic niche | Species' niche volume | 1.93 | 1.55 | 0.25 |
| | Niche overlap between primary and secondary open habitat occupied by a species | $-4.6 \times 10^{-3}$ | 0.39 | 0.91 |
| | Median of temperature niche | $-5.7 \times 10^{-2}$ | 0.13 | 0.68 |
|  | Median of precipitation niche | 0.23 | 0.25 | 0.39 |
| Functional trait | MR | -0.05 | 0.12 | 0.70 |
|  | O | -0.08 | 0.17 | 0.65 |

(A) Geographical variables:

*1. Species’ current range size* was calculated as the natural-log-transformed number of species occurrence records across all habitats.

*2. Species’ preference for open habitat* was calculated as the proportion of all occurrence records that are located in open habitat over occurrence records that are in native forests and open habitats.

*3. Availability of secondary open habitat*: In order to quantify how much open habitat has become available in the neighbourhood of primary occurrences, the availability of secondary open habitat was quantified for each species as follows: for each primary occurrence (that is, an occurrence record in forest or primary open habitat), the number of grid cells with secondary open habitat within a 10 × 10 km neighbourhood around the occurrence record was calculated. Availability of secondary open habitat is defined as the cumulative total number of secondary open grid cells. Each secondary open grid cell was not counted more than once when it was located in the neighbourhood of more than one primary occurrence.

*4. Mean elevation of current range:* to test whether species occurring at a higher elevation are more likely to take advantage of newly opened habitats, the mean elevation of all occurrence records across all habitats was calculated.

(B) Climatic variables:

*5. Species climatic niche volume:* Niche volume was estimated as a proxy of climatic tolerance and was quantified as niche overlap on 2-D space comprising of temperature and precipitation axes between each species and the entire New Zealand climate space. Niche volume was calculated using Schoener’s D index [29] with the “ecospat” package in R [30]. Schoener’s D ranges from 0 to 1 with higher values indicating larger niche overlap.

*6. Niche overlap between primary and secondary open habitat* was quantified as climatic niche overlap (Schoener’s D) between the climatic niches occupied by primary and secondary open occurrence records of each species. Higher values indicate higher similarity in climate conditions between occurrence records in primary and secondary open habitat.

*7. Medians of species temperature and precipitation niches:* The median of the temperature and precipitation axes of the species occurrences across all habitats were calculated to analyse the individual effects of temperature and precipitation on species distributions.

(C) Species trait variables:

*Life form – dispersal ability*: We selected two functional traits on the basis of relevance for the species’ ability to shift its range to analyse effects of species functional traits on species distribution. Each species was assigned one of the three combinations of life forms and dispersal ability classes; Stoloniferous-Ancistrum (eleven species), Rhizomatous-Microphyllae (five species) and other combinations (two species) (Table S3).

Based on published information [20], each species was classified as either rhizomatous (five species) or stoloniferous (eleven species) with two species belonging to other life forms. The genus *Acaena* has three distinct phylogenetic sections (Pteracaena, Ancistrum and Microphyllae; Bitter [18]) which are characterised by morphological differences in their fruits. We used these sections as an index of dispersal ability; Ancistrum has barb-tipped spines which attach the fruits to passing animals and therefore is considered as having higher dispersal ability than barbless species in the other two sections.

Nine factors from three main groups (Geographical features, climatic niche, functional traits) were tested using a multivariate generalized linear model (GLM). Coefficients, SE; standard error and p-values of each explanatory variables were derived from the GLM. The results of the GLM for “species’ life form – dispersal ability” were calculated using the comparisons with Stoloniferous-Ancistrum. MR: comparison between Stoloniferous-Ancistrum and Microphyllae_Rhizomatous and O: comparison between Stoloniferous-Ancistrum and other types of species’ life form and dispersal ability.

## Results

### Pre-human and current distribution of open habitat

#### Geographical distribution

Open habitat in the study region increased from 18.4% to 63.4% of the total land area since human arrival in the 13^th^ Century AD (Fig. 1 a). Currently, 15.3% of New Zealand’s land area is a primary open area, while approximately half (48.1%) of its current land area is secondary open habitat (Fig. 1 b). 91.0% of the primary open area and 47.7% of secondary open area are located in the South Island.

**Figure 1:**
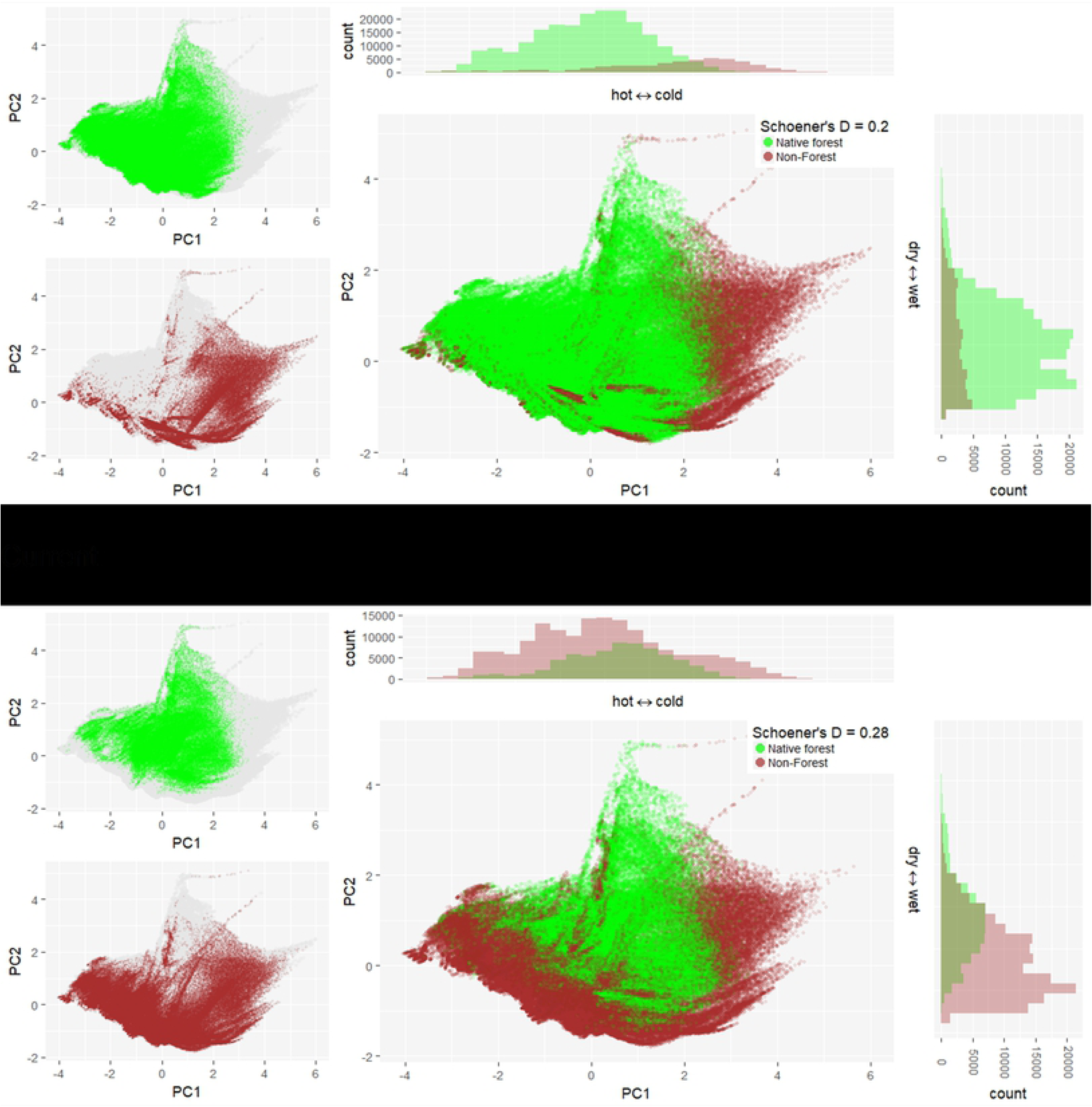
Forest and open land cover in New Zealand since human settlement at 1km grid cell resolution. (a) Forest (green) and non-forest, open (brown) land cover modelled for pre-human times in the 13^th^ century [21] and observed for current times in 2012 [22]. See Table S2 for a detailed description of land cover classes. (b) Open areas: Primary open areas (blue) indicate areas that were forest free prior to and are still open today; secondary open areas (red) are areas that were forested prior to human settlement but that are currently characterised by open habitat; others (white) area areas that are currently not open habitat or are considered unsuitable for our target species (e.g. urban area and waterbodies).

#### Climate

The climate associated with open habitats in New Zealand have shifted from cold to warm conditions since the forest clearances following human settlement, however, there was no clear directional shift along the precipitation axis (Figs. 2, 3 a). The highest frequency of current open habitat comprising both primary and secondary open habitats occurred in a warmer and wetter environment (0 – 0.5 in temperature axis and −0.5 – 0 in precipitation axis) than the pre-human open habitat (2.0 - 3.0 in temperature axis and 2.0 - −1.5 in precipitation axis) (Fig. 2). Note that temperature axis is negatively correlated with actual temperature. The more of indigenous forests in a warm environment was cleared than indigenous forests in a cold environment, as shown by the highest frequency of current forest in a colder environment (0.5 - 1.0 in temperature axis) than secondary open habitats (0 – 0.5 in temperature axis) (Fig. S1)

**Figure 2:**
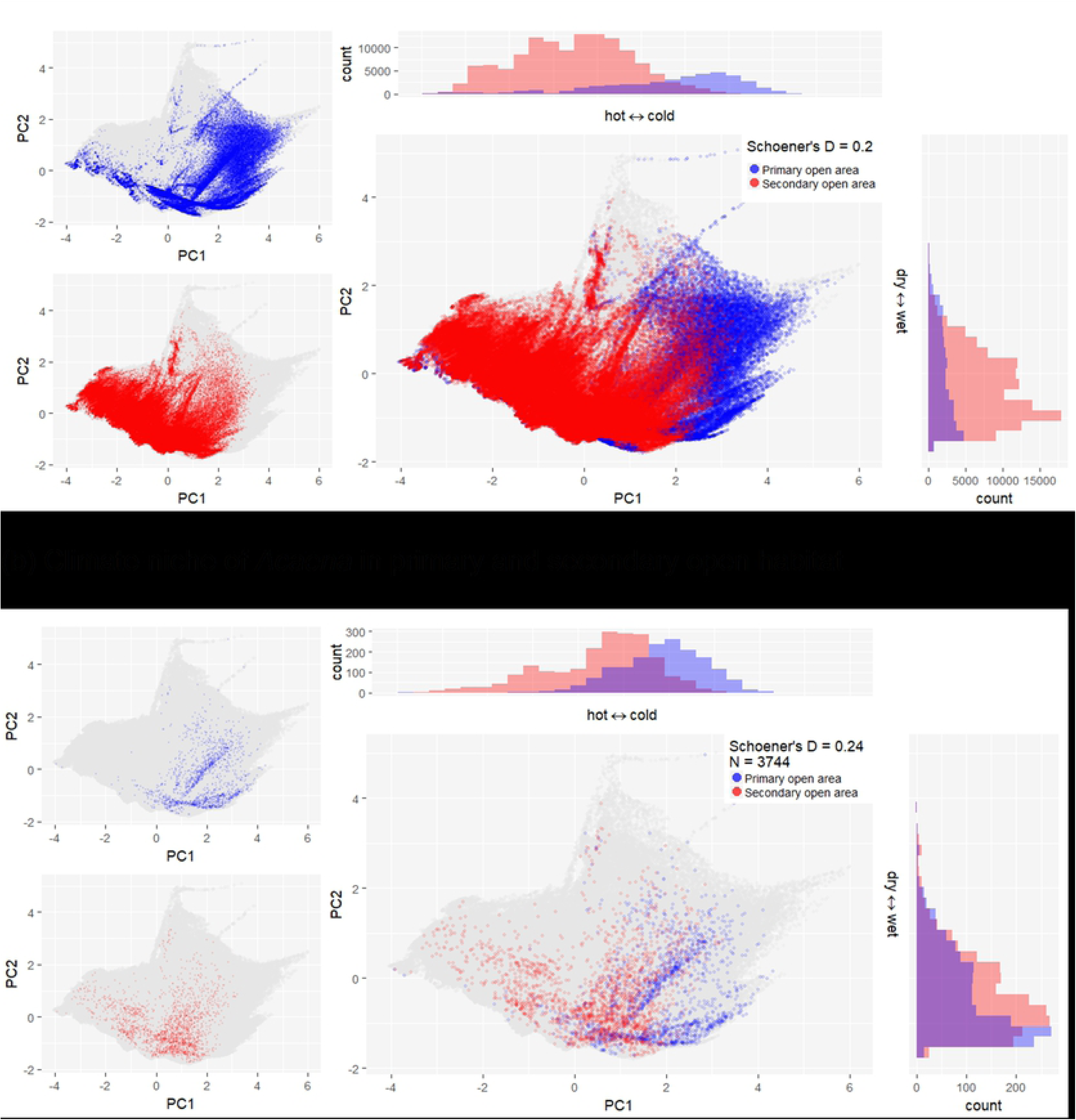
Climate conditions of forest and non-forest, open areas in New Zealand before and after human settlement. Forests are shown as green dots and open habitats are shown as brown dots. Figures show the first two axes of a Principal Component Analysis of four climate variables (see methods) at 1km grid resolution. The total climate space of New Zealand is shown in dark grey. Schoener’s D values indicate the overlap in climate conditions between forest and non-forest areas.

**Figure 3:**
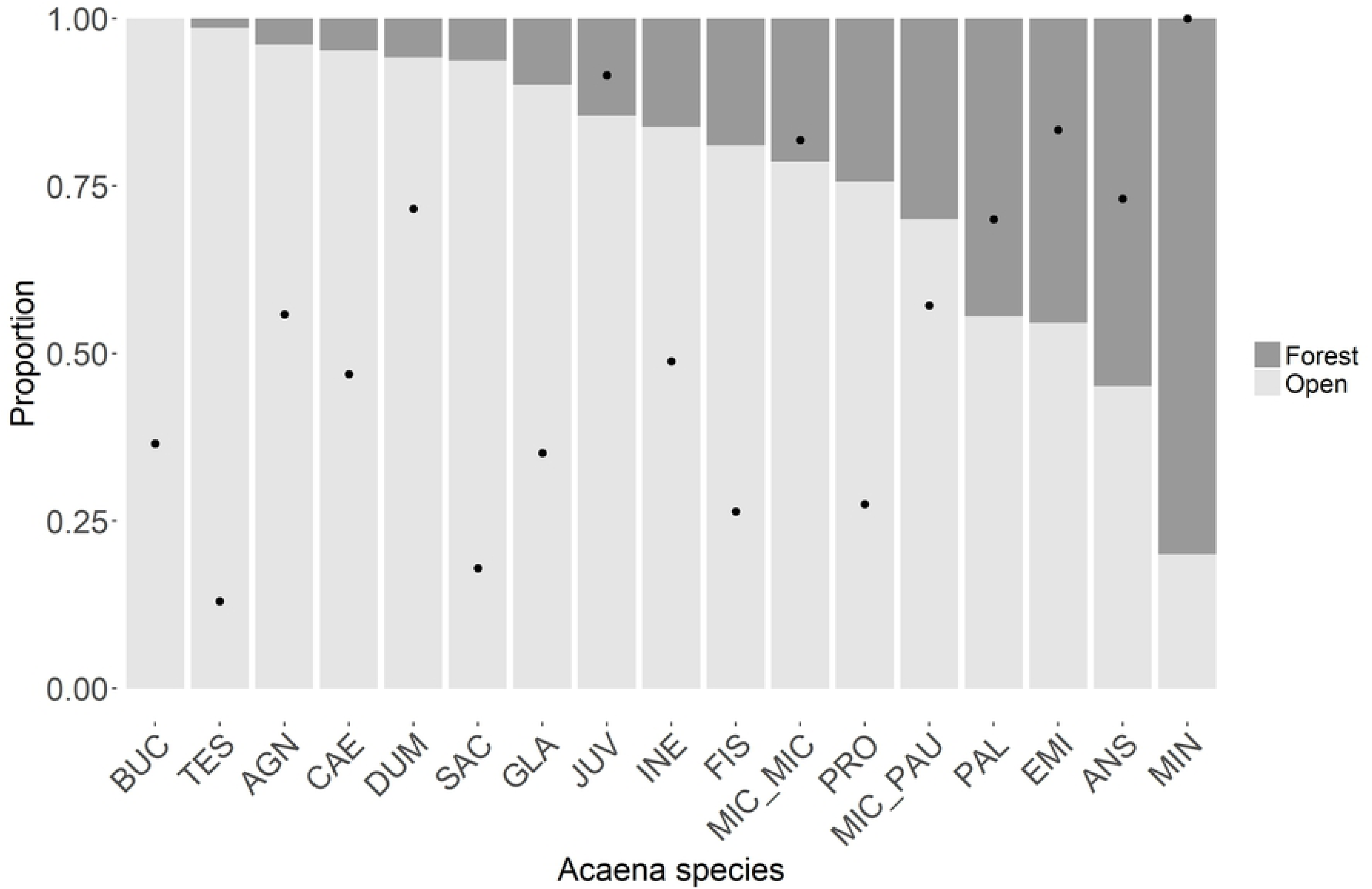
Climate space of primary and secondary open habitat areas (a) and niches of *Acaena* species in currently occupied primary and secondary open habitat (b). Schoener’s D values indicate the climate niche overlap between species’ occurrence records in primary and secondary open areas. “N” is the total number of 1 km grid cells with *Acaena* occurrences.

### *Acaena* distributions in primary vs secondary open habitat

There were 9944 occurrence records of the 18 *Acaena* species ranging from 9 to 3892 (see Fig. S2 for each species’ distribution and climatic niche). Species of *Acaena* are commonly open-habitat species, which was reflected in 68.4 % of all occurrence records of the studied species being found in currently open habitat and 26.6 % in native forest. Furthermore, the numbers of 14 species’ occurrence records in open habitats were larger than those in forests (Fig. 4). Of all occurrence records in open habitats, 46.9 % were found in primary open habitat and 53.0 % were found in secondary open habitat, indicating that *Acaena* occurrence records distribute almost equally in primary and secondary open habitats. Of all *Acaena* occurrence records, 54.9% are primary occurrences and 36.3% are secondary occurrences, given that secondary open habitat drove from the pre-human forest. The proportion of occurrence records in secondary open habitat for any species excluding *A. minor* with no occurrence records in primary open habitat ranged from 13% (*A. tesca*) to 92% (*A. juvenca*) with an average of 56% (Table S3). For eight of the 18 studied species, their proportions of secondary open habitat occupied were > 50%, indicating that they had more of occurrence records in secondary than in primary open habitat.

**Figure 4:**
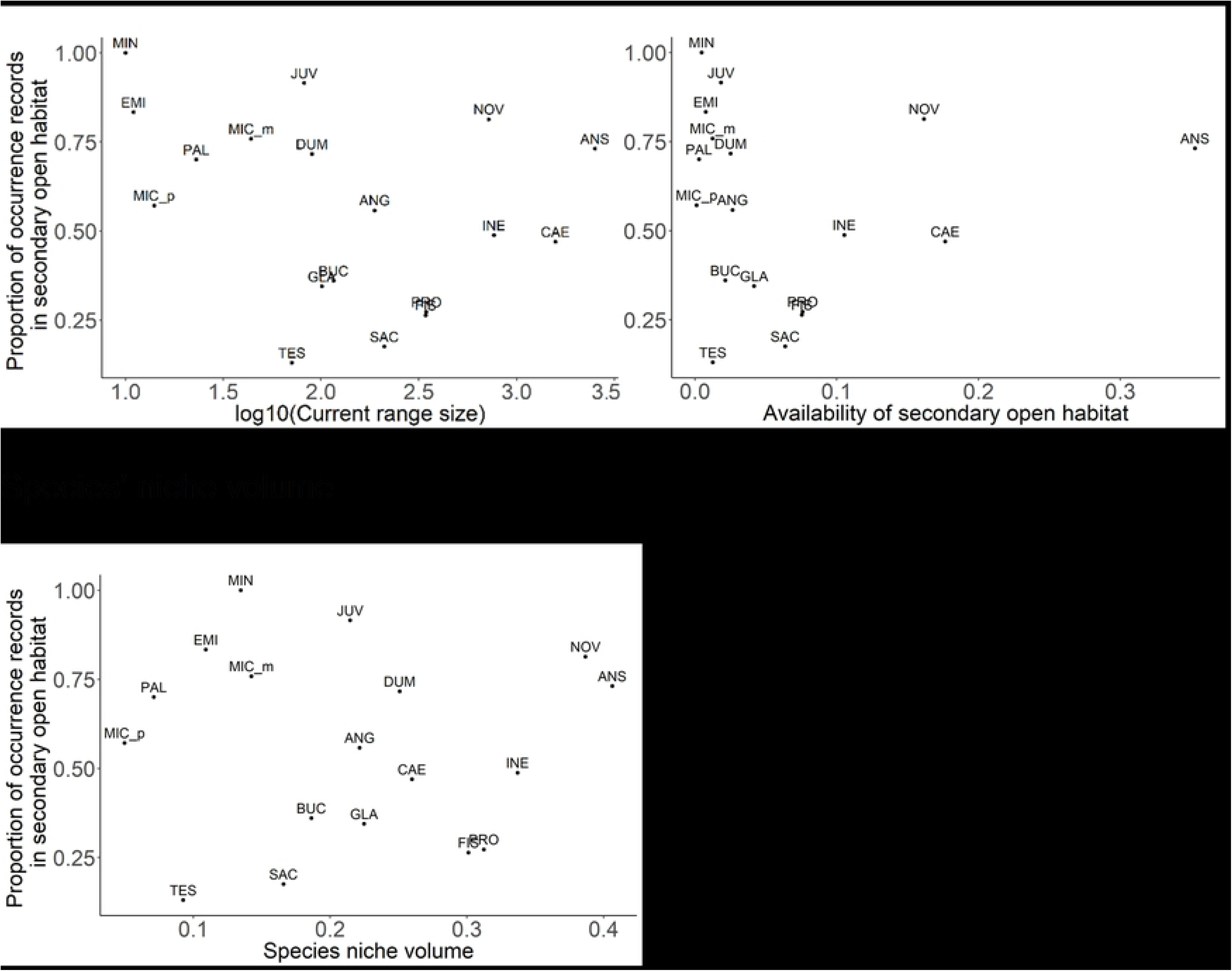
Proportion of *Acaena* species occurrence records in open (light grey) and forest (dark grey) habitats. Species are arranged in descending order of proportions of open habitat. Black dots indicate the proportion of occurrence records in secondary open habitat. See Table S1 for species name codes.

### Correlates of prevalence in secondary open habitat

#### Geography

Current range size across all habitats showed no correlation with the proportion of secondary open habitat (p = 0. 28; Fig. 5a; Table 1).

**Figure 5.**
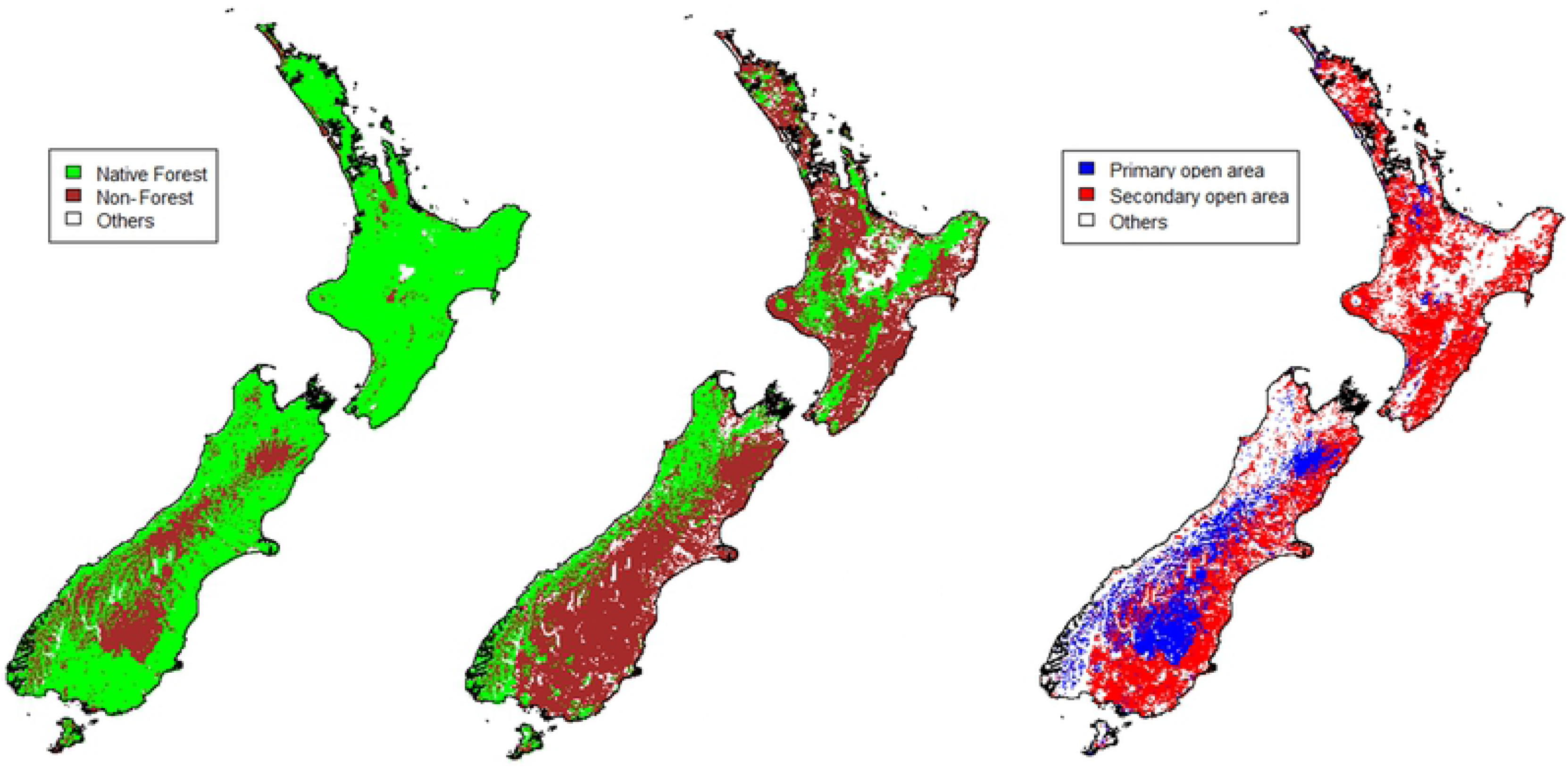
Prevalence of secondary open habitats for *Acaena* species in New Zealand and its relationship with species current range size across all habitats, availability of secondary open habitat adjacent to current *Acaena* species distribution and species’ niche volume across all habitats derived from climate parameters. See Table S1 for species name codes.

On average over the studied 18 *Acaena* species, availability of secondary open habitat was 6.6% of all secondary open area with the maximum was 35% and the minimum was 0.12%. The availability of secondary open habitat showed no correlation with proportions of secondary open habitat which species currently occupy (p = 0.43: Fig. 5b).

Preference for open habitat, the proportion of occurrence records in open habitats to forests and open habitats, ranged from 0.20 to 1 with an average of 0.77. Species preference for open habitats did not show a significant correlation with the proportion of secondary open habitat currently occupied (p = 0.39). Six species with smaller preferences for open habitats than the average over all the studied species (< 0.77) all had high proportions of secondary open habitats (> 0.5), indicating that species common in primary forests also tend to get into open habitats. The average of temperature niche medians of four strictly open habitat species, species with > 0.95 preference for open habitats, was 1.24. The average was larger than the average of temperature niche medians over all the other species (0.77), indicating that strictly open habitat species typically occur in a colder environment than the climate in which the species common in forest occur.

Over the studied 18 species, the average elevation of all occurrence records was 741 m, the maximum was 1220 m and the minimum was 38 m. Mean elevation of the species occurrence records was unrelated to the proportions of secondary open habitat (p = 0.43; Table 1). Mean elevations of species with a high preference for open habitats (> 0.75) were generally high (average; 902 m), indicating that species occurring at a high elevation generally were more likely to occur in open habitats.

#### Climate

Compared to primary open habitat, *Acaena* distributions in secondary open habitat covered larger climate spaces and showed a shift into warmer climates (Fig. 3 b). However, there was no significant relationship between the species’ niche medians on the temperature nor precipitation axes and the proportion of secondary open habitats currently occupied by the species (Median of temperature axis; p = 0.68. Median of precipitation axis; p = 0.40. Table S3).

Species climatic niche volume across all habitats ranged from 0.05 to 0.40 (mean; 0.21) (Table S3). Climate niche volume was not significantly correlated with the proportion of secondary open habitat currently occupied by the species (p = 0.25).

Over the investigated 18 species, the average niche overlap between primary and secondary open habitats was low at 0.22, indicating the climates of primary and secondary open habitats occupied by the species were generally not very similar. The maximum niche overlap between primary and secondary open habitats was 0.58 (*A. novae zelandiae*) and the minimum was 0 (*A. microphylla var. microphylla, A. saccaticupula* and *A. tesca*) (Table S3). Species niche overlap between primary and secondary open habitats was not significantly related to the proportion of secondary open habitat occupied (p = 0.91).

#### Species functional traits

There was no significant difference in proportions of secondary open habitat occupied by species among the three different types of functional traits. The mean of proportions of secondary open habitat over Stoloniferous - Ancistrum species was 62.6%, the mean over Rhizomatous - Microphyllae species was 46.2% and the mean over species with the other type was 45.1%

## Discussion

We investigated the climate conditions of pre-human and current open habitat and the prevalence of species from an open-habitat genus (*Acaena*) in secondary, i.e. recently opened habitat. We quantified the relative importance of three sets of factors – geographic features, species’ climatic niches and the species’ dispersal traits for the ability of species to utilise secondary open habitat. Our main findings are; 1) the majority of current open habitat comprising of primary and secondary open habitat is characterised by warmer climate conditions than pre-human open habitat. 2) Secondary open habitat is generally warmer than primary open habitat. 3) The prevalence in secondary open habitat varies among studied species, however, none of the measures of geographic features, climatic niche and functional traits was significant predictors of species prevalence in the deforested, secondary open habitat. Nevertheless, geographical and climatic niche factors showed stronger relationships with the species’ prevalence in secondary open habitat than functional traits associated with dispersal.

### Pre-human and current distribution of open habitats and the current distribution of open habitat plants

Since the first human settlement, c. 60% of the original, pre-human forest habitat, was transformed to open habitat in New Zealand [31]. Our results showed that the majority of current open habitats are located in low-lying warm and dry areas, in comparison with the pre-human open habitats. Pre-human open habitats were restricted to relatively small areas, mostly in the alpine areas above the natural tree line, in wetlands and riverbeds, in frosted valley floors or in dry low-lying inland areas, which generally have cold environments [32]. Low-altitude regions with warm climate were especially vulnerable to fire and are often best suited and easily accessed for agricultural conversions in New Zealand and elsewhere that deforestation happens (e.g. tropical forest [33, 34] and Latin America [35]). Therefore, open habitats in the world should have experienced the climatic shift to warmer conditions due to human activity.

### The relative importance of factors driving prevalence for secondary open habitat

#### Geography

1. Current range size across all habitats

Current range size across all habitats of *Acaena* species did not show significant correlation with the proportion of secondary open habitat occupied. Range limits can be set by climate, topography, soils and biotic interaction [36]. The factors controlling the current range limit of *Acaena* in secondary habitat are likely different from those in pre-human open habitats. It is likely that pre-human open-habitats reflected very limited climate space as they were restricted to alpine area where trees did not naturally occur [32], therefore climate of pre-human open habitat could have been insufficient for some species to realize their potential climatic niche fully, indicating that competition with forest trees was the main driver of open-habitat plant distributions in pre-human time. However, current drivers appear to vary depending on species, because the environments in secondary open habitats have broadened due to anthropogenic forest clearances, and therefore, currently available climate conditions allow open-habitat plants to obtain more of their potential climatic niche than those which they occupied before the forest clearance.

2. Availability of secondary open habitat

When new habitat becomes available for colonisation, species whose primary occurrences have more of the new habitats nearby can have advantages for expansion of their distribution into the new habitats [37]. The positive influence of historical habitat availability on grassland species richness was found in Estonian islands [38] and wood cricket populations in the UK were mainly found in woodland fragments situated closely to another occupied site [39]. However, the positive influence of habitat availability on species re-distribution was not supported by our study, in which the availability of secondary open habitat was unrelated to the proportion of secondary open habitat occupied by the species (Fig. 5b). Our method to quantify the availability of secondary open habitat did not consider possible dispersal distance (1 – 1500 m from parent plants [40]) and geographical barriers, e.g. high mountains and glacier. Glaciers worked as barriers for habitat expansion of arctic-alpine plants from the Last Glacial Maximum to date [41]. Dispersal ability is discussed further in “Species functional traits”.

3. Habitat characteristics

Characteristics of current habitats can explain prevalence in specific habitats. Although *Acaena* species are generally open-habitat species, some species can occur within forests and in edge habitats between forests and open habitat (e.g. *A. anserinifolia*) [17]. In terms of their current distribution, species with a high preference for open habitats seem to be restricted in more open habitats (e.g. grasslands), while species with a low preference for open habitats tend to occur in less open habitat (e.g. shrublands) frequently (Fig. S3). Both grasslands and shrublands were considered open habitats in our study, however, they have different levels of openness. Species with a low preference for open habitat should be shade-tolerant, indicating that the species would survive in less open habitat.

In general, species occurring mainly at higher elevations occupy smaller areas of secondary open habitat, which was indicated by the negative relationship between means of elevation of current range and proportions of secondary open habitat (Table 1). This appears to represent specialisation to colder conditions, and therefore, indicates more restrictions on the species expansion into secondary open habitats. For instance, species whose primary habitat was restricted to the alpine/montane area and/or colder regions showed very small proportions of secondary open habitat (e.g. *A. saccaticupula* and *A. tesca*).

### Environmental space

Species with larger climatic niche volumes did not have significantly greater occupancy in secondary open habitats (Table 1). This result is against the idea that niche breadths predict geographical range size [42]. However, temperature niches of *Acaena* species were generally a better predictor of species geographical range expansion than precipitation. Species that mostly occur in cold primary open area (> 0 of temperature axis) tend to occupy a small proportion of secondary open habitat (e.g. *A. saccaticupula* and *A. tesca*). Deforestation in New Zealand expanded substantial open habitats in warmer climates, however, had small impacts on extending the availability of these habitats across rainfall gradients.

### Species functional traits

Functional traits associated with regeneration and dispersal are critical for establishing populations in new habitats [40]. Laanisto, Sammul [43] showed a strong relationship between distribution change and functional traits across 736 species in diverse genera. Barb-spined *Acaena* species (species in Ancistrum section) generally showed broad geographical ranges and habitat distributions (Fig. S2) and have higher adherence to animals than barb-less species [44, 45]. However, life form and dispersal ability of *Acaena* did not show any relationships with species prevalence of secondary habitats. This result supports Lloyd, Lee [46] showing no consistent trait differences between common and rare species and could be attributed to far greater dispersal efficiency following the human arrival with the introduction of many small mammals, stock, particularly sheep and cattle, and granivorous birds [47]. The difference of dispersal ability tested in our study was just an improvement of an adhesive feature of seeds to animals, which does not change dispersal types. In addition, the frequent occurrences of *Acaena* beside roads and tracks reported by Lloyd, Lee [46] indicate that human transport has established novel pathways for the spread of *Acaena,* as well as for alien species all over the world [48].

### Mechanism of realized niche change

Some of *Acaena* species have obtained new climatic niche with obtaining secondary open habitat. There are two possible mechanisms of how *Acaena* obtained new climatic niche:

1) Niche evolution; species of *Acaena* from forest and primary open habitats adapted to the new habitats created by deforestation, expanding environmental tolerance and range.

2) Competitive release; species of *Acaena* were released from the competition with forest plant species, which triggered the expansion of *Acaena* distribution into newly opened habitats.

#### 1. Niche evolution

Niche conservatism constrains the environmental expansion and diversification of species because of inherent restrictions in adaptive plasticity [49]. However, niche evolution along specific climatic parameters is poorly understood. The evolution rates of potential environmental niches appear variable. Petitpierre, Kueffer [50] showed that change of climatic niche through emigration into new habitats is uncommon for terrestrial plants and Wasof, Lenoir [51] showed that niche can be conserved up to 10^4^ years. On the other hand, Early and Sax [52] showed that a large proportion of species’ naturalized distributions occurred outside the climatic conditions occupied in their native ranges.

#### 2. Competitive release

Change of land cover can have drastic and rapid effects on species distribution. Realized environmental niche can change on the change of non-climatic factors [e.g. species traits and land use; 53, 54, 55], if they play a strong role in limiting species’ native distributions. Land cover change can destroy habitats of some species, however, it can let other species to shift their habitat into new habitats by releasing them from competitions. Species expansion through release from competition has been found in animals where a reduced predator community contributed to modern fishers’ range expansion [56]. The competitive release is a more realistic mechanism for the change in species prevalence in open habitat than the evolution of *Acaena* species’ climatic niche, because evolutionary processes of adaptions to new environments generally take a long time.

Although some species traits can change in a shorter period (e.g. change of timing of phenological events as the reaction to climate change [57]), evolutionary change of species traits (e.g. morphological change) generally requires significantly more time. Therefore, the time since when *Acaena* species have obtained their new climatic niche (c. 800 years) appears too short for them to evolve their climatic niches.

The response of *Acaena* species to new habitats and climates in our study suggests that species will exhibit varying distributional shifts as the climate warms. The most vulnerable *Acaena* species would be those restricted to colder montane/alpine habitats, where physiological specialisation can restrict options in a warming world. This indicates that global warming could lead to habitat loss and elevational range shifts of species restricted to colder montane/alpine habitats as the upward elevational range shifts which have been reported globally [58, 59].

### Limitations

Our study is based on the land cover data on 1 km grid resolution which can be too coarse scale for measuring ecological processes. Moreover, a single land cover class was assigned to each grid, homogenising any fine-scale habitat diversity. Plant response to environmental change can take decades or longer [60]. However, it is likely that the herbaceous studied species have attained equilibrium with the new climate regime after 800 years from the human arrival, because this time frame is much longer than lags in climate response reported in other studies [e.g. 40, 59, 61, 62]

### Conclusions

Land cover change is a key component of global environmental change driving the redistribution of species as a consequence of human activity. Change from closed forest to open habitat is a typical feature of anthropogenic environmental change providing new and more area suitable for open-habitat species. The climate conditions of current open habitats in New Zealand are warmer than pre-human times, because forests in warm lowland were cleared for hunting and agricultural purposes. Thus, anthropogenic activity has opened new parts of the available climate space for open-habitat species. Our result suggested that open-habitat species could have occupied only parts of their potential climate space at the LGM, because they were kept out of climatically suitable areas through competition with trees. Whereas at the present, open-habitat species possibly occupy larger parts of their climatically-suitable areas than those at the LGM because these areas have been made tree-free by humans. We found that overall geographical and environmental factors were more important than species functional traits for potentially facilitating expansion into secondary habitats.

## Acknowledgements

We thank J.B. Steel for species observation data and occurrence records.

## Supporting information

**Figure S1. Climate space of forest (green) and secondary open habitats (blue) in New Zealand after human settlement.** All the figures show climate space of New Zealand as darker grey background. Climate space of native forests (top left) and open habitat (bottom left) areas are shown respectively. The centre figures show climate space of the two habitats together. The histograms on the right side show numbers of 1 km grid cells along temperature (top) and precipitation (right) axes.

**Figure S2. Maps and climate space of primary (blue) and secondary (red) occurrences of *Acaena* species.** “N” in the legend of maps shows the number of occurrences in primary and secondary open area. The total climate space of New Zealand is shown in dark grey.

**Figure S3. Proportion of *Acaena* species occurrences in LCDB land cover classes and proportion of secondary open habitat.** LCDB land cover classes were coloured by a habitat type and levels of openness; open habitat with low openness (blue gradient colours), open habitat with high openness (yellow gradient colours) and forests (green gradient colours). Black points on bars show species’ proportion of secondary open habitat. Proportion of secondary open habitat for *A.minor* (”MIN” in the figure) is 1 due to no occurrence records in primary open habitat. Bars were sorted in descending order of preference for open habitat. Species name codes are shown in Table S1.

**Table S1. List of species and their number of occurrence records and habitats.**

**Table S2. List of land cover classes in original pre-human and current land cover data and classes after conversion to 1 km grid cell data.**

**Table S3. List of analyzed variables; proportion of secondary open habitat and 9 environmental predictors.** SO: Proportion of secondary open habitat, the number of secondary open occurrence records divided by the number of primary and secondary open occurrence records. Niche overlap; values of Schoener’s D showing climate niche overlap between primary open occurrence records and secondary open occurrence records

Author contributions
WGL, RO and BJA conceived the study, MN developed and conducted the analyses and wrote the first draft of the manuscript. All authors contributed to the writing and editing of the manuscript.

